# RAD18 Methylation by the Methyltransferase SETD6 Attenuates DNA Breaks

**DOI:** 10.1101/2025.06.27.662082

**Authors:** Lital Estrella Weil, Michal Feldman, Jennifer Van Duine, Ji Qiu, Joshua LaBaer, Dan Levy

## Abstract

This study investigated the interaction between the SETD6 lysine methyltransferase and RAD18, a key protein in the DNA damage repair pathway. SETD6 belongs to the SET-domain-containing family of proteins, which are known to catalyze protein methylation, a post-translational modification that plays a critical role in regulating protein function, stability, and interactions. Using protein microarray technology, we identified RAD18 as an interactor and substrate of SETD6. We confirmed this interaction through ELISA and immunoprecipitation assays, demonstrating that SETD6 directly binds and methylates RAD18. Using mass spectrometry and site-directed mutagenesis, we identified that RAD18 undergoes mono-methylation at the K73 and K406 residues. Furthermore, we found that RAD18 methylation affects its nuclear localization. Specifically, SETD6 KO cells exhibited increased nuclear RAD18 levels, suggesting that methylation status influences RAD18’s shuttling between the cytoplasm and nucleus. Notably, depletion of SETD6 led to elevated markers of DNA damage (γH2AX) and increased DNA breaks, as evidenced by comet assays. Restoring SETD6 activity significantly reduced DNA damage, while a catalytic inactive mutant did not have this effect, underscoring the importance of SETD6’s enzymatic function. Overall, our results demonstrate that SETD6-mediated methylation of RAD18 is essential for attenuating DNA breaks, thereby regulating its cellular localization and function in maintaining genomic integrity.

## Introduction

Protein lysine methylation is catalyzed by a large class of enzymes known as lysine methyltransferases (KMTs), of which there are over 60 candidate members. Currently, the majority of KMTscontain a conserved catalytic Su(var)3-9, Enhancer-of-zeste, Trithorax (SET) domain, which catalyzes the addition of methyl groups to histone and non-histone proteins to thereby regulate diverse cellular processes (1–4). SETD6 is a mono-methyltransferase which regulates numerous cellular, biological and physiological processes such as the NF-κB cascade (5–7), the NRF2 oxidative stress response (8), WNT signaling (9, 10) and nuclear hormone receptor signalling (11).

RAD18 is a RING-finger E3 ubiquitin ligase which is involved in the DNA damage response (DDR) pathway. Within the DDR pathway, the canonical role of RAD18 is catalyzing the monoubiquitination of PCNA (12). RAD18 is essential in the DDR pathway, particularly in post-replication repair pathways. It is recruited to double strand break (DSB) sites forming foci which are co-localized with 53BP1, NBS1, phosphorylated ATM, BRCA1 and γH2AX. In addition, RAD18 was shown to promote DSB repair during G1 phase through chromatin retention of 53BP1 (NHEJ)(13) .

Additionally, cells carrying a double mutation, one in RAD18 and the other in a gene that functions in homologous recombination (HR) or non-homologous end joining (NHEJ), were found to be synthetically lethal. Increasing evidence indicates that the expression levels of RAD18 play a critical role in mutagenesis and the response to DNA damage(14–16). This highlights the importance of understanding the regulatory mechanisms that govern RAD18’s cellular functions.

In a proteomic screen using a novel protein microarray technology (Multiplexed Nucleic Acid Programmable Protein Array (M-NAPPA)(17), we identified, among other proteins, RAD18 as a potential interacting partner of SETD6. Biochemical and cellular experiments revealed that SETD6 binds and methylates RAD18 at K73 and K406. Mechanistically, our findings reveal that the methylation status of RAD18 governs its selective recruitment to the nucleus, which in turn influences the levels of γH2AX and the extent of DNA breaks. This suggests that the regulation of RAD18 by SETD6 could play an important role in the cellular response to DNA damage.

## Results

### SETD6 binds and methylates RAD18

In order to identify SETD6 interactors and potential substrates, we utilized a novel protein microarray technology (Multiplexed Nucleic Acid Programmable Protein Array (M-NAPPA) **(Fig. 1A)**, which enables high throughput detection of protein-protein interactions (17). Seventy-seven potential SETD6 interactors were identified from this screening (**Fig. S1**). Among these, RAD18 was identified and we sought to examine a direct interaction between SETD6 and RAD18 using purified recombinant SETD6 and RAD18 proteins. The direct interaction between SETD6 and RAD18 was assessed with an ELISA assay, where a direct interaction between GST-SETD6 and His-RAD18 was observed (**Fig. 1B**). BSA and GST served as negative controls. The homodimerization of SETD6 (18) served as positive control for these experiments. To further test the interaction of SETD6 and RAD18 in cells, we performed an Immunoprecipitation (IP) assay. As shown in **Figure 1C**, SETD6 interacts with immunoprecipitated RAD18. These findings raised the possibility that RAD18 can serve as a substrate to SETD6. To address this hypothesis, we performed an *in-vitro* radioactive methylation reaction in the presence of recombinant His-SETD6, His-RAD18 and ^3^H-SAM (*S*-adenosyl-methionine) as the methyl donor. We found that SETD6 methylates RAD18 (**Fig. 2A**). In addition, we found that the methylation of immunoprecipitated FLAG-GFP-RAD18 from PC3 cells increases in the presence of recombinant SETD6 (**Fig. 2B**). To validate whether SETD6 methylates RAD18 in-cells, we immunoprecipitated methylated cellular proteins from control and SETD6 KO cells using a pan-Methyl antibody. We found that the methylation of RAD18 in HeLa cells was decreased in the SETD6 knockout cells (**Fig. 2C**). Since SETD6 KO cells do not display a complete abolishment of RAD18 methylation, we assume that RAD18 might be methylated by additional methyltransferases.

**Figure 1.**
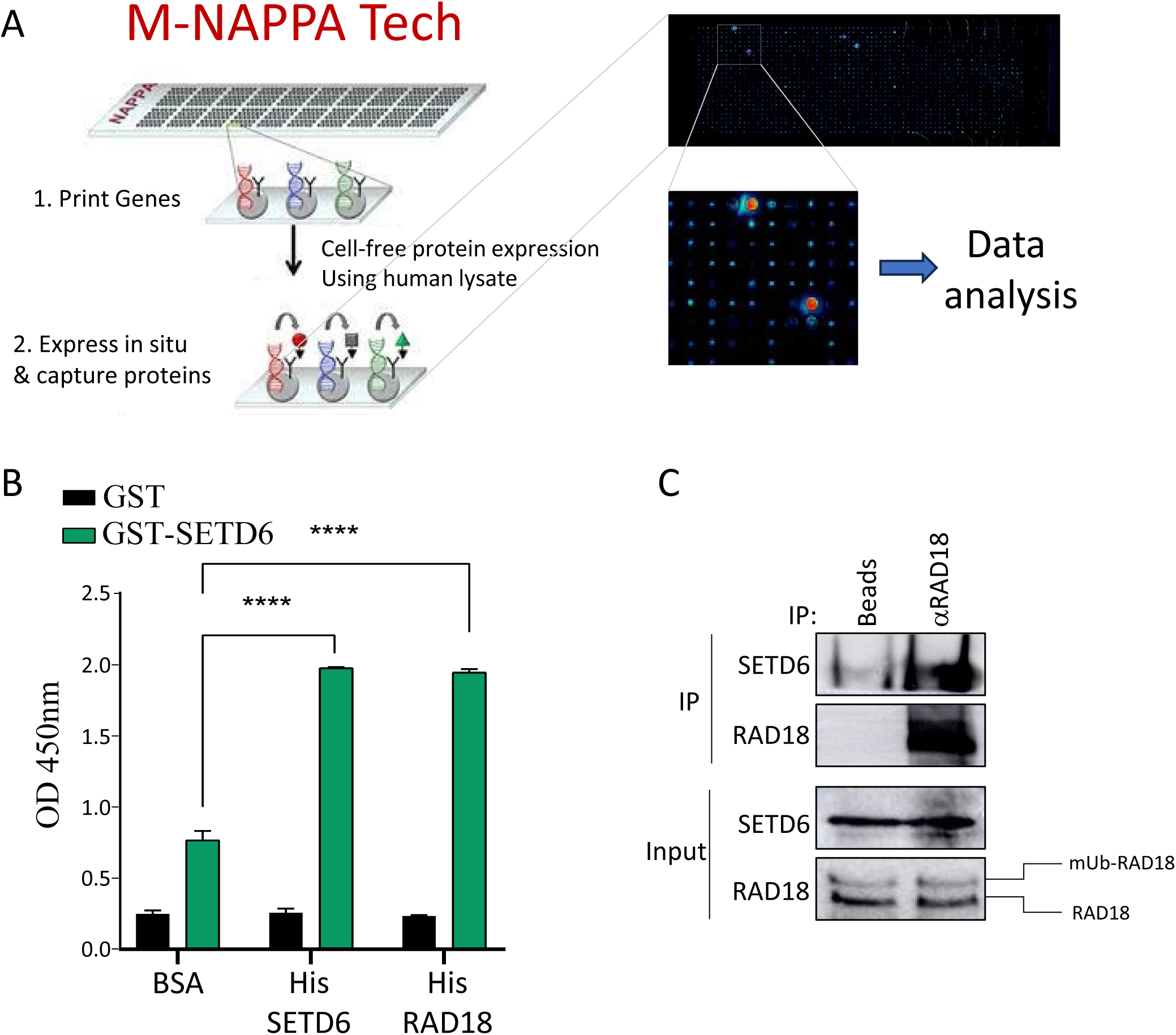
SETD6 binds RAD18 in-vitro and in cells. **(A)** Schematic representation of the M-NAPPA process. DNA sequences encoding specific proteins were spotted onto a glass slide. The DNA is then expressed *in-situ* using a cell-free expression system, creating a protein microarray. The interaction was detected using fluorescence following data analysis. **(B)** Enzyme-linked immunosorbent assay (ELISA) was performed with the indicated recombinant proteins. Self-interaction between His-SETD6 and GST-SETD6 served as a positive control. The graph represents absorbance at 450nm for each condition. ****p-value <0.0001**. (C)** HeLa whole cell lysates were immunoprecipitated with agarose beads conjugated to the indicated antibody, followed by Western blot analysis with indicated antibodies.

**Figure 2.**
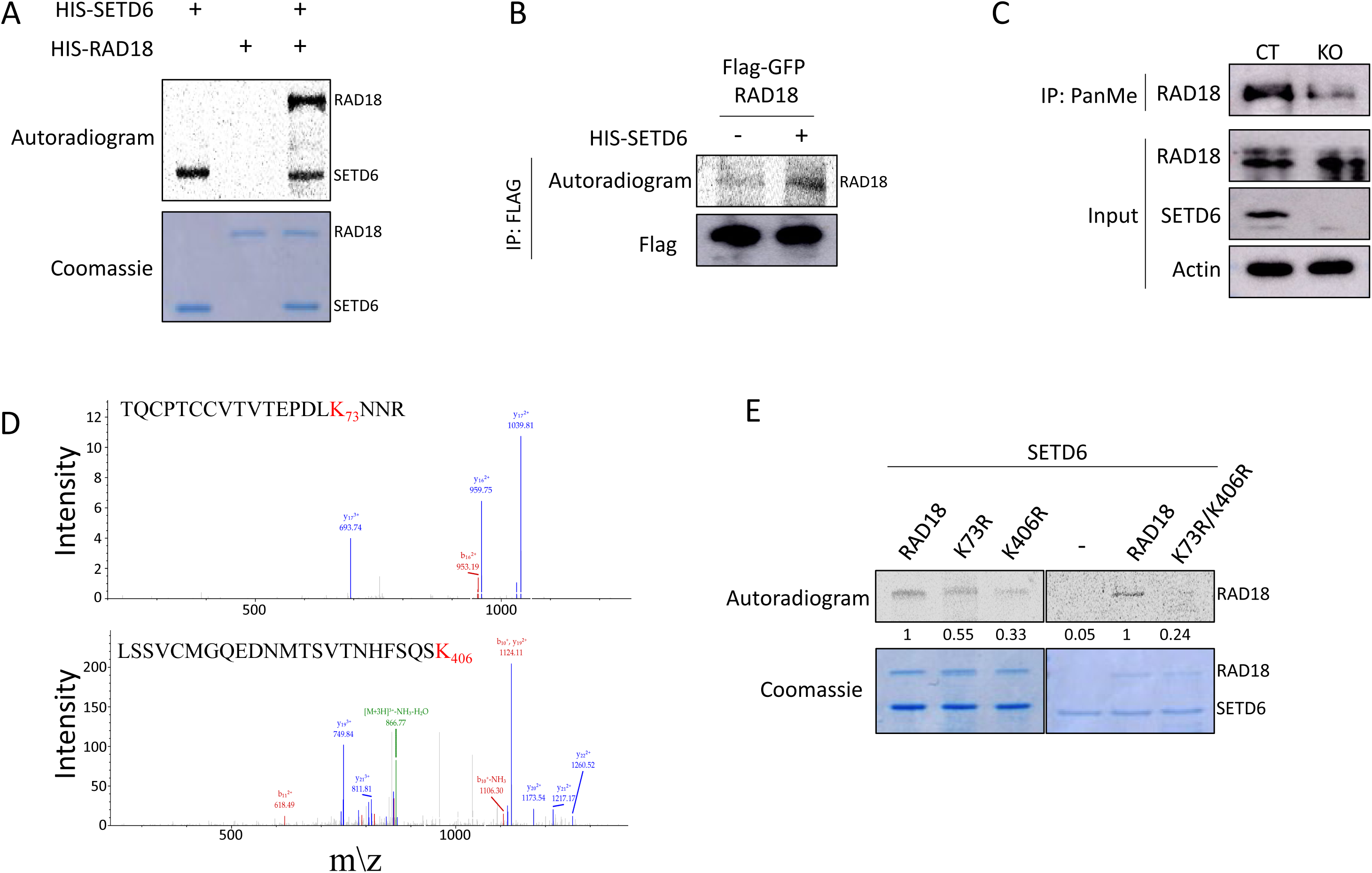
*SETD6* methylates RAD18 on K73 and K406. **(A)** *In-vitro* methylation assay in the presence of ^3^H-labeled SAM with recombinant His-SETD6 and His-RAD18. The methylated proteins were detected by autoradiogram and a Coomassie stain of the recombinant proteins used in the reactions is shown on the bottom. **(B)** Semi-*in-vitro* methylation assay of immunoprecipitated Flag-RAD18 that was over-expressed *in PC3 cells,* with and without the presence of His-SETD6. The methylated proteins were detected by autoradiogram and Flag-RAD18 was visualized using Western blot analysis using αFlag antibody. **(C)** HeLa cells lysates were immunoprecipitated with Pan-Methyl lysine antibody followed by Western Blot analysis with the indicated antibodies. **(D)** MS spectra of RAD18 TQCPTCCVTVTEPDLK73NNR and LSSVCMGQEDNMTSVTNHFSQSK406. Major y- and ions are displayed as red, and blue, respectively. **(E)** *In-vitro* methylation assay in the presence of ^3^H-labeled SAM with the indicated recombinant proteins. Coomassie stain of the recombinant proteins used in the reactions is shown on the bottom.

To map RAD18 methylation site(s), recombinant His-RAD18 was incubated under methylation reaction conditions with and without His-SETD6, followed by Mass spectrometry (MS) (**Fig. 2D**). Two potential mono-methylation sites were identified: K73 and K406. Interestingly, both residues are not conserved in mouse and yeast (**Fig. S2**). To validate the MS findings, we have generated individual methylation-deficient RAD18 lysine-to-arginine mutants at K73 and K406 (K73R and K406R) as well as a double mutant (K73R/K406R) using site-directed mutagenesis. Using a radioactive methylation assay, we observed a decrease in the methylation signal for both individual mutants, while it was almost abolished in the double mutant (**Fig. 2E**). These results revealed that K73 and K406 are the primary methylation sites on RAD18 by SETD6.

### RAD18 methylation restricts its nuclear accumulation

RAD18 has a dynamic cellular shuttling pattern between cytosol and the nucleus (19), thus we hypothesized that SETD6 might affect RAD18 accumulation to the nucleus. To examine this hypothesis, we utilized immunofluorescence to detect endogenous RAD18 in control and SETD6 KO cells **(Fig. 3A and WB in Fig S3A)**. To evaluate the total integrated density of RAD18 signal in the nucleus, cells were visualized using confocal microscopy (LSM) and analysis was performed using the Fiji program. The results indicate a higher level of RAD18 in the nucleus in KO cells compared with control cells. To test if this phenomenon depends on the methylation status of RAD18, we repeated this experiment in cells expressing either Flag-RAD18 WT or the methylation-deficient Flag-RAD18 K73R/K406R mutant **(Fig. 3B)**. We observed a significant higher amount of nuclear Flag-RAD18 K73R/K406R mutant compared to Flag-RAD18 WT. These results demonstrate that nuclear RAD18 levels are negatively regulated bySETD6 as well asRAD18 K73/K406 methylation.

**Figure 3.**
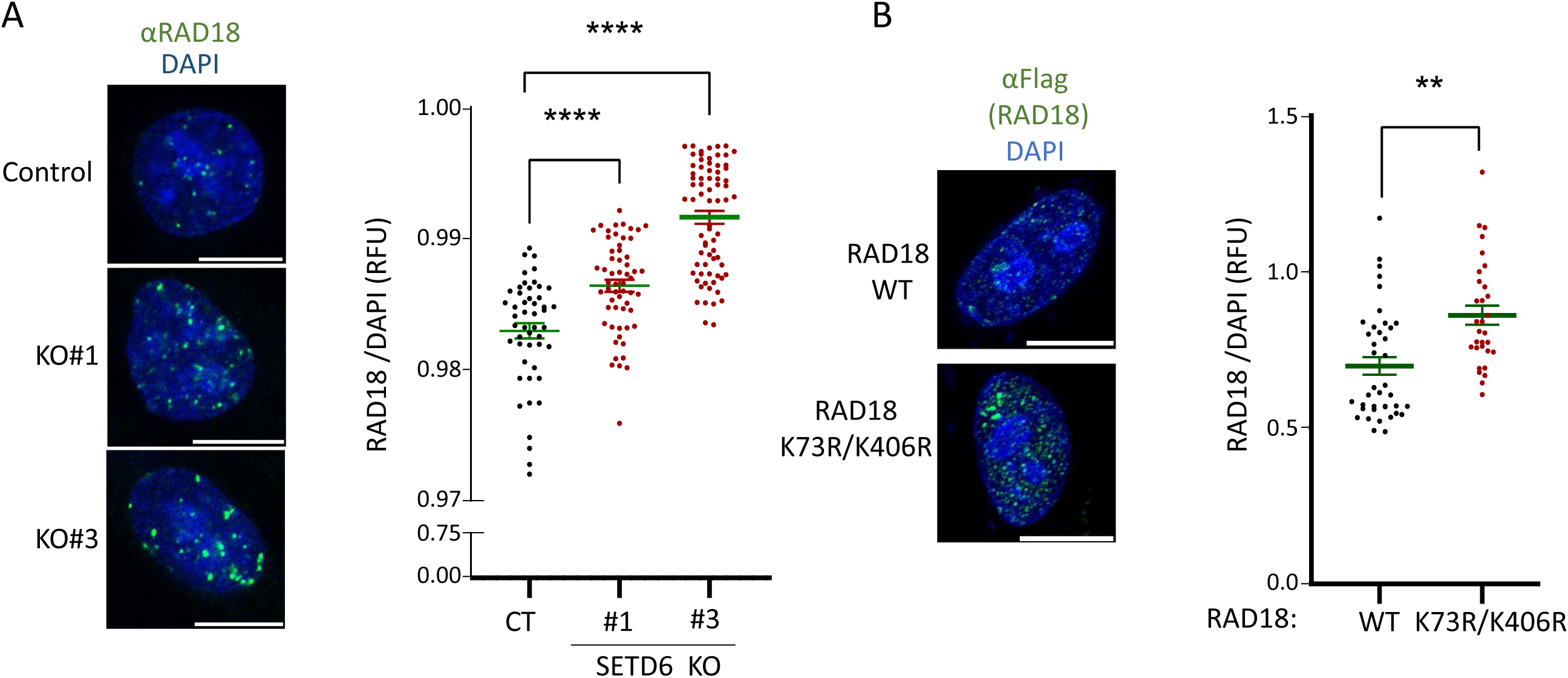
RAD18 is differentially recruited to the nucleus based on its methylation state. **(A)** Control and SETD6 KO cells were stained for DNA (DAPI) and endogenous RAD18, as indicated. Total integrated density of RAD18 in the nuclei was analyzed using Fiji software. **(B)** HeLa cells overexpressing Flag-RAD18 WT or K73R/K406R were stained for DNA (DAPI) and Flag, as indicated. Total integrated density was analyzed as in A. Graph values represent mean ± SEM. **p-value <0.01 ****p-value <0.0001. Scale, 10μM.

### RAD18 methylation is linked to DNA damage accumulation

Our initial immunofluorescence experiments presented in Figure 3 show that RAD18 displays punctate staining pattern in the nucleus which was also reported before (20, 21). This observation suggested that RAD18’s recruitment to the nucleus may be linked to DNA damage accumulation. Interestingly, induction of γH2AX level by doxorubicin treatment in control cells is mimicked by SETD6 KO **(Fig. 4A)**. Consistent with these results, we observed significantly higher levels of γH2AX in SETD6 KO cells in immunofluorescence experiments **(Fig. 4B)**. These results demonstrate that SETD6 restrains the DNA damage response under basal cellular conditions. To further assess these observations, we performed a Neutral Comet Assay (Comet) which measures DSBs (22). In agreement with **Fig. 4A** which shows low levels of γH2AX in control cells, the comet assay did not reveal the presence of DNA breaks in control cells (**Fig. 4C**). However, a higher tail DNA percentage rate was observed in SETD6 KO cells. To further confirm whether this process depends on SETD6 enzymatic activity, we introduced Flag-SETD6 WT or the catalytically-dead Y285A mutant (6)into the SETD6 KO cells. While WT SETD6 significantly decreased the rates of accumulated DNA breaks, the catalytic inactive mutant failed to do so (**WB in Fig. 4D and Fig. 4E)**. Finaly, our results suggest that this phenomenon is RAD18 methylation dependent as overexpression of Flag-RAD18 K73R/K406R displayed significant higher DNA breaks compare to Flag-RAD18 WT expressing cells (**Figure 4F, and WB in Fig S3B**). Thus, RAD18 K73/K406 methylation by SETD6 restrains DNA breaks and maintains genomic integrity under basal cellular conditions.

**Figure 4.**
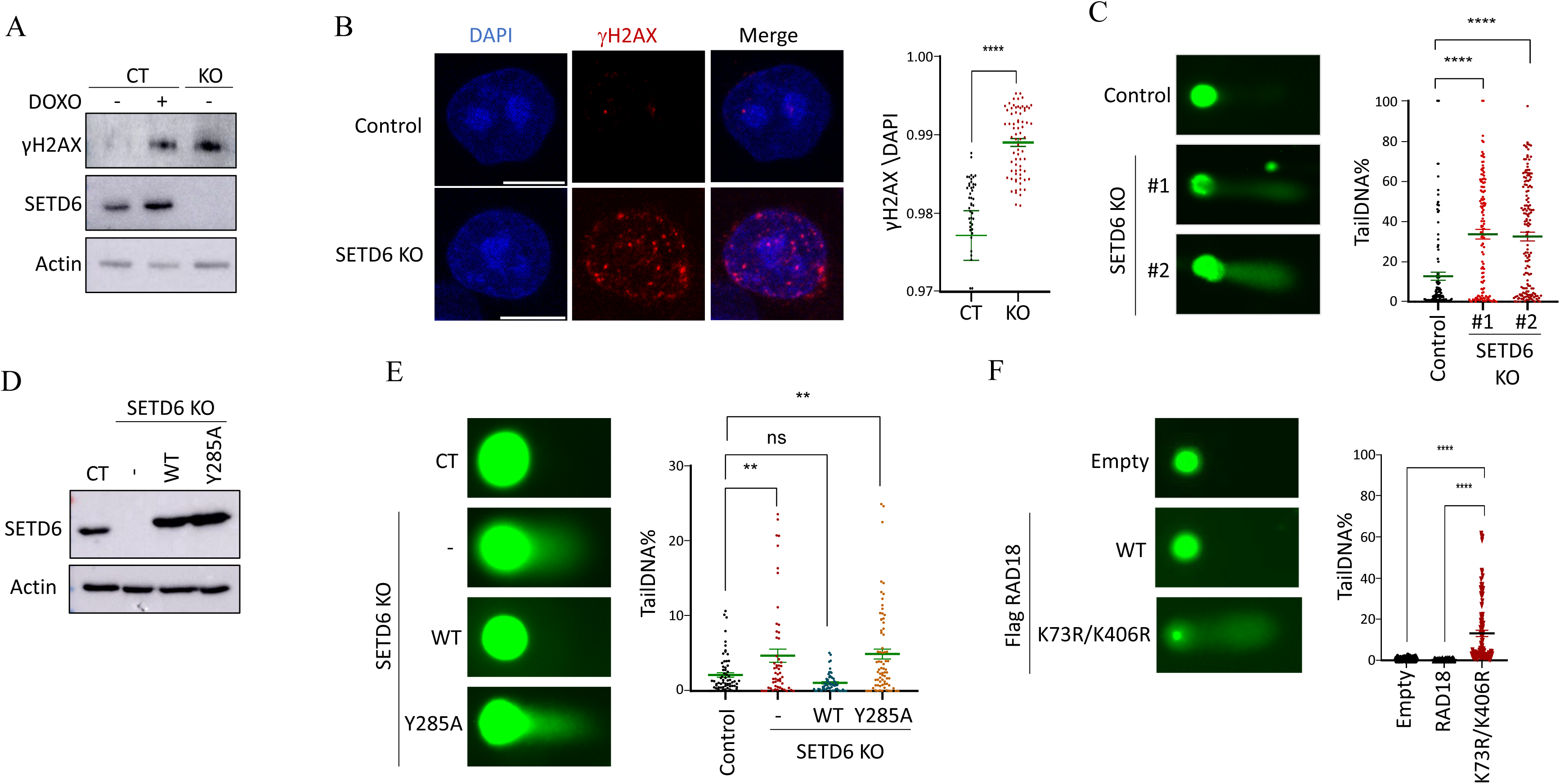
RAD18 methylation is linked to DNA damage accumulation. **(A)** Control and SETD6 KO cells were subjected to Western blot analysis to evaluate γH2AX levels. Control cells pre-treated with Doxorubicin served as positive control. Actin served as loading control. **(B)** Control and SETD6 KO HeLa cells were stained for endogenous γH2AX, as indicated. Total integrated density of γH2AX was analyzed using Fiji software. Graph values represent mean ± SEM. ****p-value <0.0001. **(C)** Neutral comet assay in control and SETD6 KO cells is presented with representative Images taken with Evos fluorescence microscope and analysis was performed using CaspLab software. Graph values represent mean tail DNA%. **(D)** Western blot analysis with the indicated antibodies for CT and SETD6 KO cells rescued with SETD6 WT or SETD6 Y285A catalytic inactive mutant **(E)** HeLa control, SETD6 KO and SETD6 KO cells rescued with Flag-SETD6 and SETD6 Y285A catalytic inactive mutant and were treated as in C. Graph values represent mean ± SEM. *p-value <0.05 **p-value <0.01 ****p-value <0.0001. Scale, 10μM. **(F)** Cells overexpressing RAD18 WT or K73R,K406R were subjected to Neutral comet assay as in C. Graph values represent mean ± SEM ****p-value <0.0001. Scale, 10μM.

## Discussion

Earlier studies have established SETD6 to be a key regulator of various biological processes including inflammation, oxidative stress, cell cycle regulation, and cellular differentiation (4). In this study, we reveal a previously unknown role for SETD6 in DNA-damage response. Using biochemical and cellular approaches, we provide evidence that SETD6 interacts and methylates RAD18 at K73 and K406. Our data supports a model by which the methylation of RAD18 at K73 and K406 by SETD6 enhances RAD18 recruitment to the nucleus. Knockout of SETD6 results in higher levels of γH2AX and DNA breaks. Our data further suggests that this phenomenon depends on the enzymatic activity of SETD6 and RAD18 methylation at K73 and K406.

RAD18 undergoes multiple post-translational modifications at several residues (23–25). Further, RAD18 is known to physically interact with SETD1A, another protein lysine methyltransferase (26). However, our study provides the first evidence demonstrating that RAD18 undergoes lysine methylation. We show that RAD18 methylation was decreased in SETD6 KO cells, indicating that RAD18 is methylated by SETD6. However, RAD18 methylation was not completely abolished in the absence of SETD6, suggesting that RAD18 might be methylated by an additional methyltransferase at other residues. It is not uncommon that proteins involved in DNA damage response are targeted for methylation by multiple methyltransferases to regulate their activity. For example, following DNA damage, p53 can undergo methylation at lysine 372 by SET9, which enhances the stability of chromatin-bound p53 (27). Conversely, SET8 methylation at lysine 382 suppresses p53 transcription activity (28). Raising a specific antibody targeting K73/K406 methylation is required to assess the contribution of these residues to RAD18 cellular processes.

Interestingly, K73, which was identified as a target for SETD6 methylation, is also predicted to be a monoubiquitinated site (29, 30). A recent paper has shown that RAD18 is phosphorylated at S403 by ATR (31), which is adjacent to K406 which is methylated by SETD6. K406 phosphorylation disrupts the interaction between Rad18 and PCNA. Thus, it will be interesting to examine potential crosstalks between the different modifications that can may orchestrate the regulation of RAD18 cellular function.

The K73R/K406R mutant showed a more dramatic decrease in methylation *in-vitro* compared to each individual mutation; K73R or K406R. Thus, we chose to conduct phenotypic experiments using the double mutant. However, further studies should also examine each individual mutant, as each one may possess a unique influence on RAD18 cellular activity. Given that K73 resides in proximity to the RING domain (32), it would be interesting to examine its effect on RAD18 self-dimerization and as a result, its mono-ubiquitination and localization to the nucleus. On the other hand, K406 resides at the Polƞ binding domain (33) and may indicate that the methylation on K406 might affect RAD18-Polƞ interaction and, in turn, be involved in the regulation of the TLS pathway.

In conclusion, our study highlights the regulatory role of SETD6 in restraining DNA breaks under basal conditions through the methylation of RAD18. While our research concentrated on RAD18 activity under basal conditions, we believe that further investigation into the precise mechanisms and functional consequences of SETD6-dependent RAD18 methylation during DNA damage will enhance our understanding of DNA damage response and repair processes.

## Materials and Methods

### Plasmids and primers

For recombinant purification, SETD6 and RAD18 sequences were amplified by PCR and subcloned into pET-Duet plasmids. RAD18 mutants were generated using site-directed mutagenesis and cloned into pET-Duet. Primers sequences are presented in Table 1)

**Table 1.**
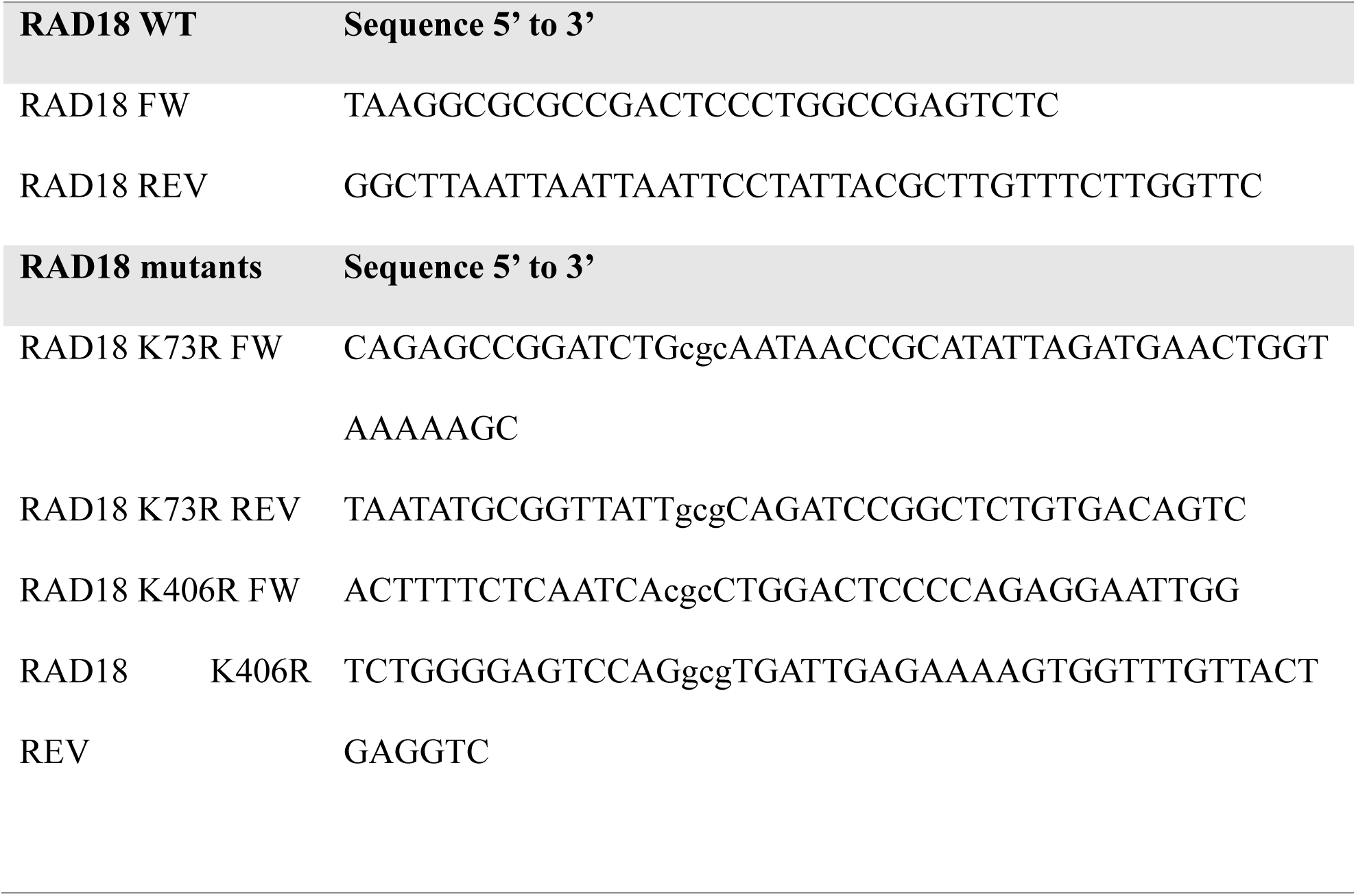
Primers for cloning and mutagenesis.

### Cell Lines, Transfection, Infection and Treatment

HeLa and PC3 cell lines (ATCC) were maintained in Dulbecco’s modified Eagle’s medium (Sigma, D5671) with 10% fetal bovine serum (FBS) (Gibco), penicillin-streptomycin (Sigma, P0781), 2mg/ml L-glutamine (Sigma, G7513) and non-essential amino acids (Sigma, M7145), at 37°C in a humidified incubator with 5% CO2. For cell transfection, cells were transfected using Mirus TransIT®-LT1 reagent, according to the manufacturer’s instructions. Transfection was performed for 24-48h before harvesting the cells.

### Antibodies, Western Blot Analysis and Immunoprecipitation

Primary antibodies used were: anti-FLAG (Sigma, F1804), anti-Actin (Abcam, ab3280), anti-anti-GAPDH (Abcam, G041), anti-Pan-methyl (Abcam, ab23366), anti-SETD6 (Genetex, GTX629891), anti-RAD18 (Cell Signaling, #9040), anti GST (Abcam, ab9085), anti-His (Thermo Fisher scientific, rd230540a) and anti-gH2AX (Merck, 05-636-25UG, clone JBW301).

HRP-conjugated secondary antibodies (goat anti-rabbit and goat anti-mouse) were purchased from Jackson ImmunoResearch (111-035-144 and 115-035-062, respectively). Fluorescently labeled secondary antibodies used: Alexa488 anti-mouse (Invitrogen, R37120) and Alexa488 anti-rabbit (Invitrogen, A21441).

For Western blot analysis, cells were homogenized and lysed in RIPA buffer (50 mM Tris-HCl pH 8, 150 mM NaCl, 1% Nonidet P-40, 0.5% sodium deoxycholate, 0.1% SDS, 1 mM DTT, and 1:100 protease inhibitor mixture (Sigma)). Samples were resolved on SDS-PAGE, followed by Western blot analysis.

Immunoprecipitation was performed using protein A/G agarose beads (Santa Cruz, SC-2003) conjugated to the antibody of interest. Briefly, ∼200μg of proteins extracted from cells using RIPA buffer were incubated overnight at 4°C with 15μl pre-washed FLAG M2 beads or protein A/G beads. The beads were then washed 3 times with RIPA buffer, heated for 5 min in Laemmli sample buffer at 95°C and resolved on a 7%-10% SDS-PAGE gel followed by Western blot analysis.

### Recombinant Protein Expression and Purification

pET-Duet containing SETD6 or RAD18, WT or mutants, with 6xHis-tag, were transformed into *Escherichia coli*, Rosetta. After IPTG induction, the bacteria were grown in LB medium at 18°C overnight (SETD6) or 37°C for 6h (RAD18). A cell pellet was isolated from medium using centrifugation (10 min, 7,000 rpm, 4 °C) and lysed by sonication on ice for 3 min in total, 10/20 on/off, 45% amplitude. Finally, the lysate was centrifuged (20 min, 18,000 rpm, 4°C) and filtered (0.22μM), and the His-tagged proteins were purified on a HisTrap column (GE) with the ÄKTA system. Proteins were eluted by 0.5M Imidazole followed by dialysis to 10% glycerol in PBS buffer. GST-SETD6 was expressed and purified as previously described (34).

### Enzyme-Linked Immunosorbent Assay (ELISA)

For the SETD6-RAD18 interaction test, approximately 2ug of His-RAD18 or His-SETD6 diluted in PBS were added to a 96-well plate (Greiner Microlon) and incubated for 1h at room temperature followed by blocking with 3% BSA for 30 min. Then, the plate was covered with 0.5μg GST-SETD6 or GST protein (negative control) diluted in 1% BSA in PBST for 1h at room temperature. Plates were then washed and incubated with primary antibody (anti-GST, 1:4000 dilution) followed by incubation with HRP-conjugated secondary antibody (goat anti-rabbit, 1:2000 dilution) for 1h. Finally, TMB reagent followed by 1N H2SO4 (stop solution) were added; absorbance at 450nm was detected using the Tecan Infinite M200 plate reader.

### M-NAPPA analysis

M-NAPPA analysis to identify SETD6’s substrates was performed as described in Yu & Song *et.al*., 2017 (17). Briefly, DNA sequences encoding specific proteins were spotted onto a glass slide. The DNA is then expressed in-situ using a cell-free expression system, creating a protein microarray. The interaction was detected using fluorescence following data analysis. The cDNA for SETD6 was inserted into pJFT7-nHalo vector and those for candidate binding proteins were inserted into pANT7-cGST vector. For ELISA validation, we first coated the 96-well plate with 50µl 1:500 anti-goat-GST. SETD6 and candidate proteins were then co-expressed with the Hela cell lysate-based IVTT system (Thermo Fisher) *in-vitro*, and the expressed proteins were diluted 1:200 in milk-PBST and added into the coated 96-well plates. The SETD6-candidate protein complex was captured by the anti-GST antibody at the bottom of the well and HRP-conjugated anti-Halo antibody was then added to the well. The presence of the interaction is indicated by a positive HRP-TMB reaction measured at an absorbance at 450nm. FOS-nHalo and JUN-cGST were used as positive controls, and SETD6-nHalo and GST only as negative controls.

### *In-Vitro* Methylation Assay

Methylation assay reactions contained 1μg of His-RAD18 WT or mutant and 4μg of His SETD6; 2mCi ^3^H-labeled S-adenosyl-methionine (SAM) (Perkin-Elmer, AdoMet); and a PKMT buffer (20 mM Tris-HCl pH 8, 10% glycerol, 20 mM KCl, 5 mM MgCl2). The reaction tubes were incubated overnight at 30°C. The reactions were resolved by SDS-PAGE for Coomassie staining (Expedeon, InstantBlue) or autoradiography.

### Semi *In-Vitro* Methylation Assay

Cells were transfected with FLAG-RAD18 plasmid. Cells were lysed using RIPA lysis buffer as described above. The samples were then washed 3 times with PBS buffer and once with PKMT buffer, followed by an *in-vitro* radioactive methylation assay overnight at 30°C, in the presence of 1μg His-SETD6. The reactions were resolved by SDS-PAGE for Coomassie staining or autoradiography.

### CRISPR/Cas9-Knock-Out

For CRISPR/Cas9-mediated gene disruption, two different sgRNAs for SETD6 were cloned to the plentiCRISPR plasmid (Addgene #49535) and sequence validated as described in Chen *et al.* [29]. Following transfection and puromycin selection, single clones were isolated and expanded as described in (34).

### Comet Assay

Cells were grown in 6-well culture plates at 5×10^5^ cells/well for 24 hours. Then, Comet Assay was conducted on cells under physiological conditions or cells that were subjected to DNA damage induction with 0.001% MMS treatment for 1h followed by an over-night recovery, as indicated. The DNA damage accumulation was determined by applying the Comet Assay Kit (3-well slides; ab238544) according to the manufacturer’s instructions. The DNA damage was expressed as tail DNA percent and tail length, calculated using the CaspLab software.

### Immunofluorescence and Image processing

For immunofluorescence (IF), cells were fixed with 4% PFA or methanol and permeabilized with 0.5% Triton X-100 and blocked with 10% fetal bovine serum for 30 min. Then, cells were stained for 3h with the relevant primary antibody followed by a 30 min incubation with fluorescent secondary antibody and DAPI. For fluorescence quantification, images were acquired with a confocal laser scanning microscope ZEISS LSM880 confocal microscope using the Plan Apochromat 63x/1.4 oil DIC M27 objective.

Image processing and analysis was performed using Fiji. For the quantification of fluorescence intensity, the nuclear area was segmented using the wand tracing tool in and integral fluorescence intensity for corresponding channel was calculated within the defined area. Data was normalized to the integral fluorescence density of DAPI staining obtained in the same area.

### Mass Spectrometry

Samples of non-radioactive methylation assay containing 2μg His-RAD18 and 4μg His-SETD6 were incubated with SAM overnight at 30°C. An additional sample without SAM served as reference. Samples were then submitted to MALDI-T**O**F analysis at The Nano Institute, BGU.

### Bioinformatic and Data Analysis

Multiple Sequence Alignment (MSA) of orthologous RAD18 methylation sites was conducted using the Clustal Omega tool (https://www.ebi.ac.uk/Tools/msa/clustalo/).

### Statistical analyses

Statistical analyses for all assays were performed with GraphPad Prism software, using one-way, two-way analysis of variance (ANOVA) or the student’s t-test.

## Acknowledgments

We thank the Levy and LaBaer labs for technical assistance and helpful discussions. This work was supported by grants to DL from The Israel Science Foundation (262/18 and 496/23), The Research Career Development Award from the Israel Cancer Research Fund and from the Israel Cancer Association.

Lital Estrella Weil^1^, Michal Feldman^1^, Jennifer Van Duine^2^, Ji Qiu^2^, Joshua LaBaer^2^ and Dan Levy^1#^

## Author Contribution

LEW, and DL conceived and designed the majority of the experiments. JVD, JQ and JL performed the NAPPA screening experiment. MF helped with the microscopic data analysis and provided valuable conceptual input for the study. LEW and DL wrote the paper. All authors read and approved the final version of the manuscript.

## Data availability

The datasets used and/or analysed during the current study available from the corresponding author on reasonable request.

---

**Figure S1.**
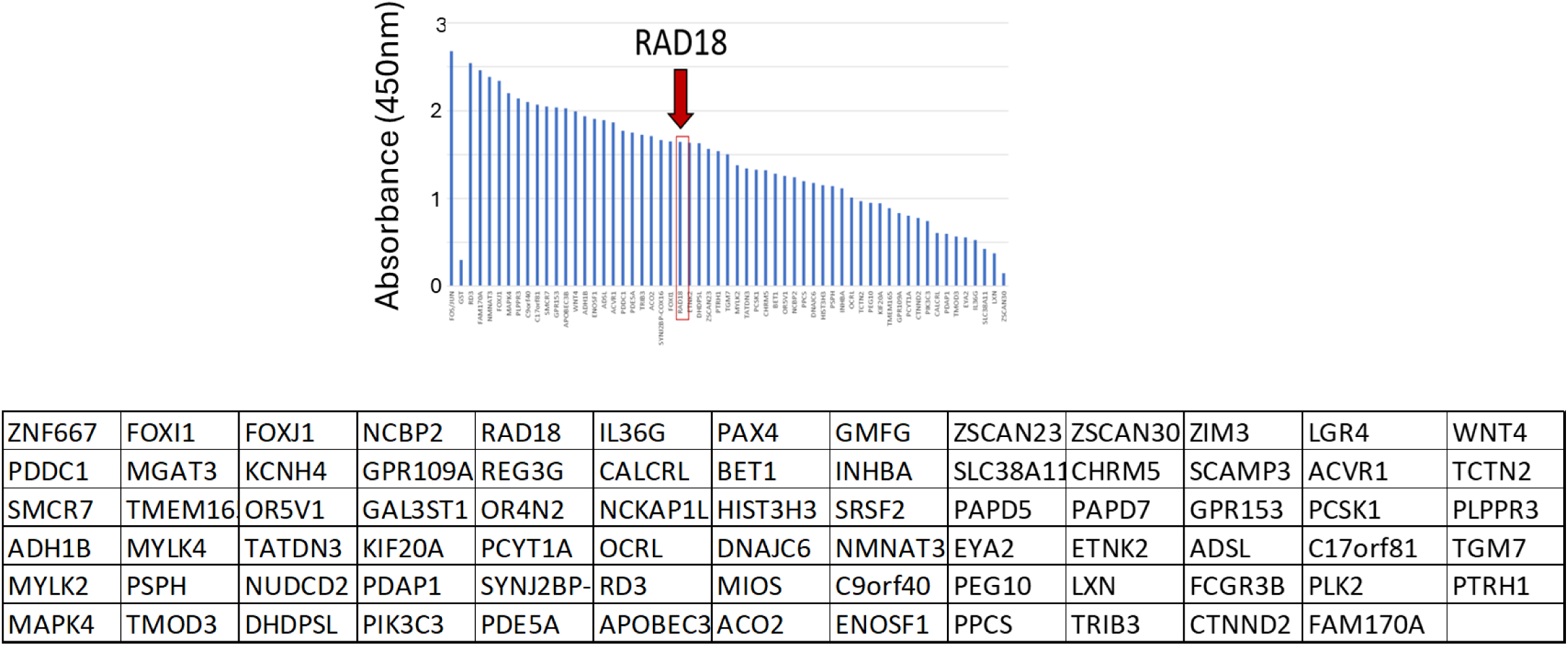
ELISA assay was conducted to detect interactions between SETD6 and candidate proteins that were then co-expressed with the Hela cell lysate-based IVTT system (Thermo Fisher) *in-vitro*. FOS-nHalo and JUN-cGST were used as positive controls, and SETD6-nHalo and GST only as negative controls. The interaction was detected by a positive HRP-TMB reaction measured at an absorbance at 450nm.

**Figure S2.**
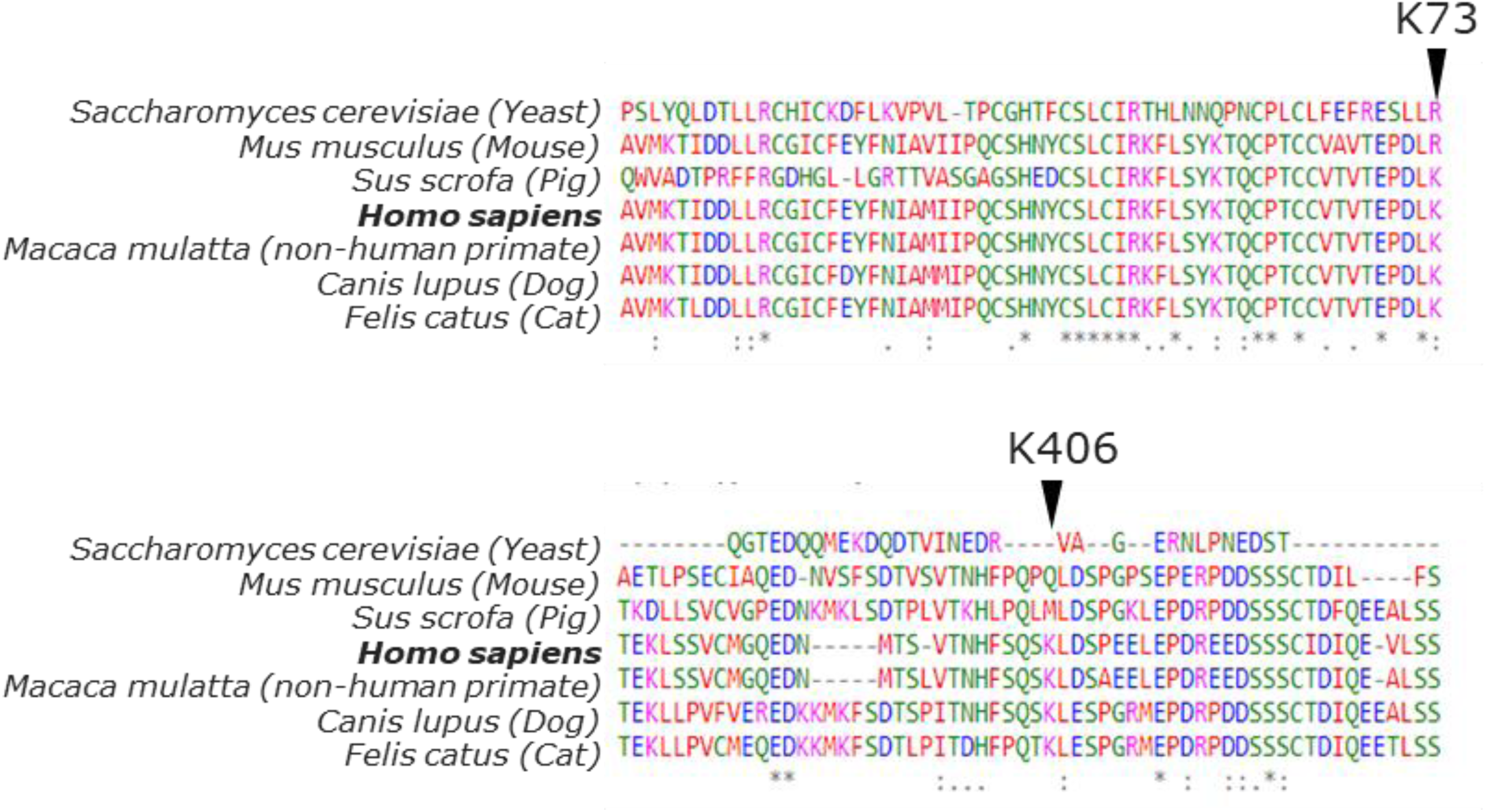
A multiple protein sequence alignment of RAD18 K73 and K406 potential methylation sites. Multiple protein sequence alignment (MSA) conducted using Clustal Omega tool for the indicated orthologous organisms.

**Figure S3.**
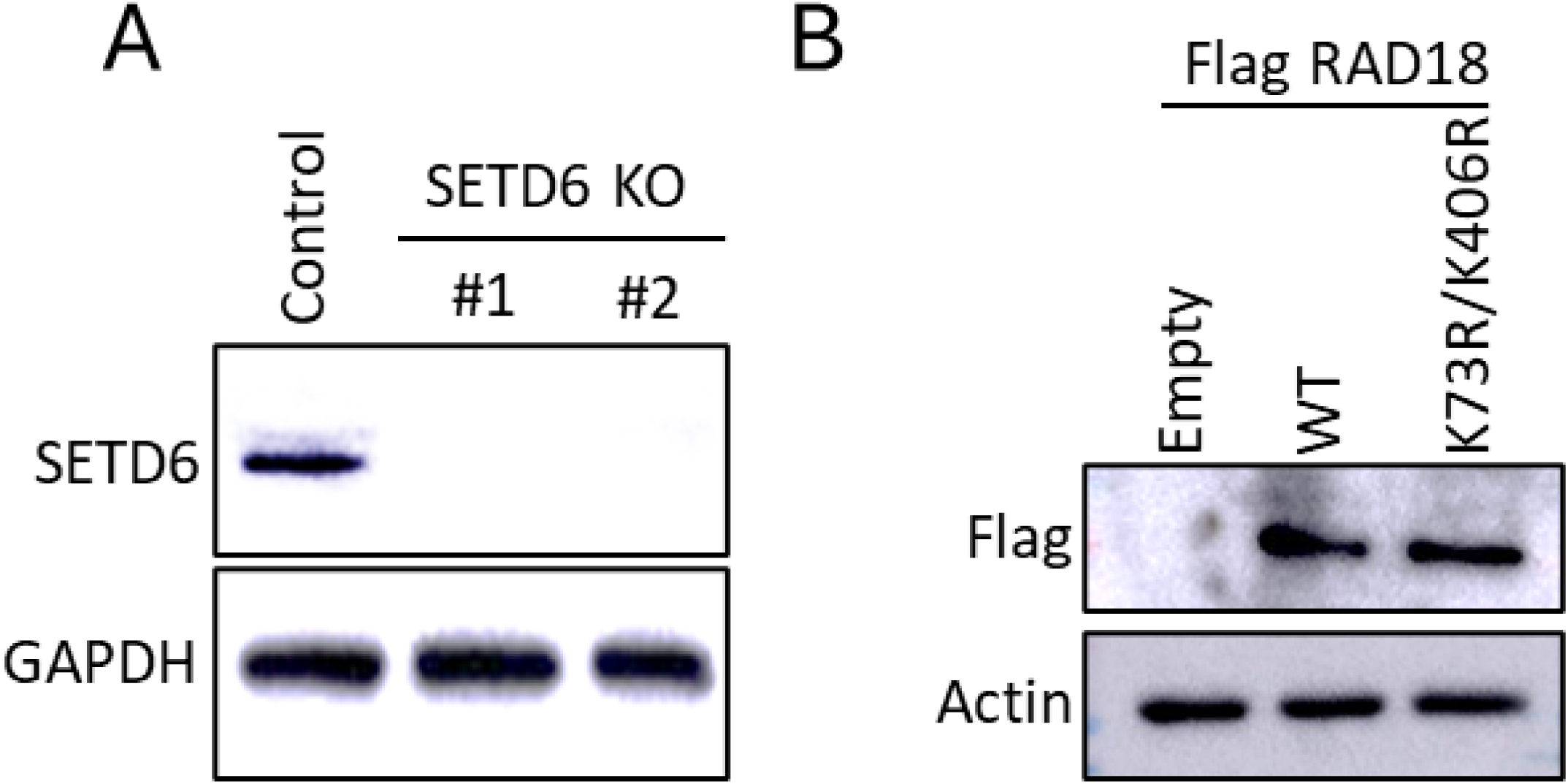
(A+B) Western blot analysis with the indicated antibodies for CT and SETD6 cells (A) and for cells expressing Flag-RAD18 WT and K73R/K406R (C).

**Fig S4.**
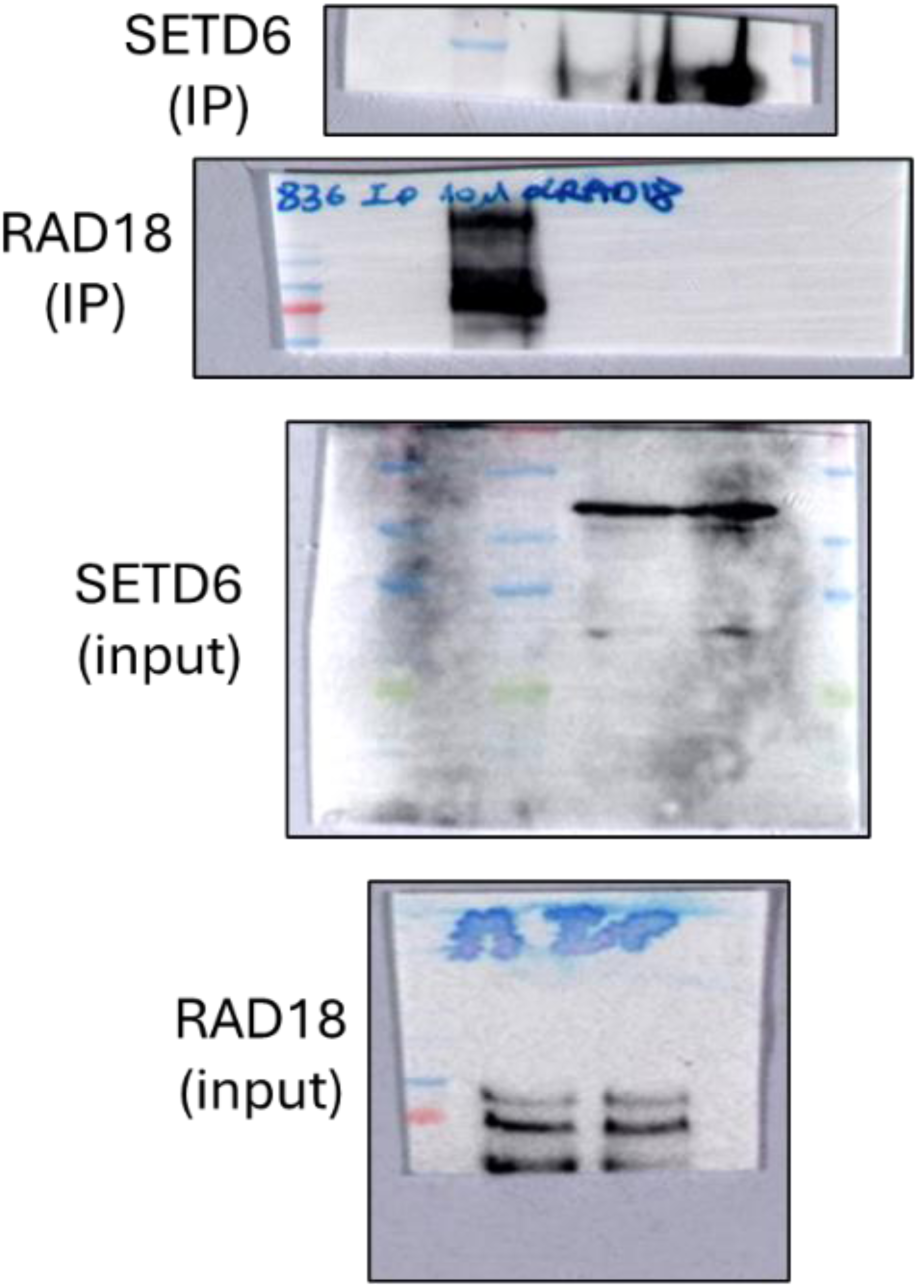
Original blots for Figure 1C

**Fig S5.**
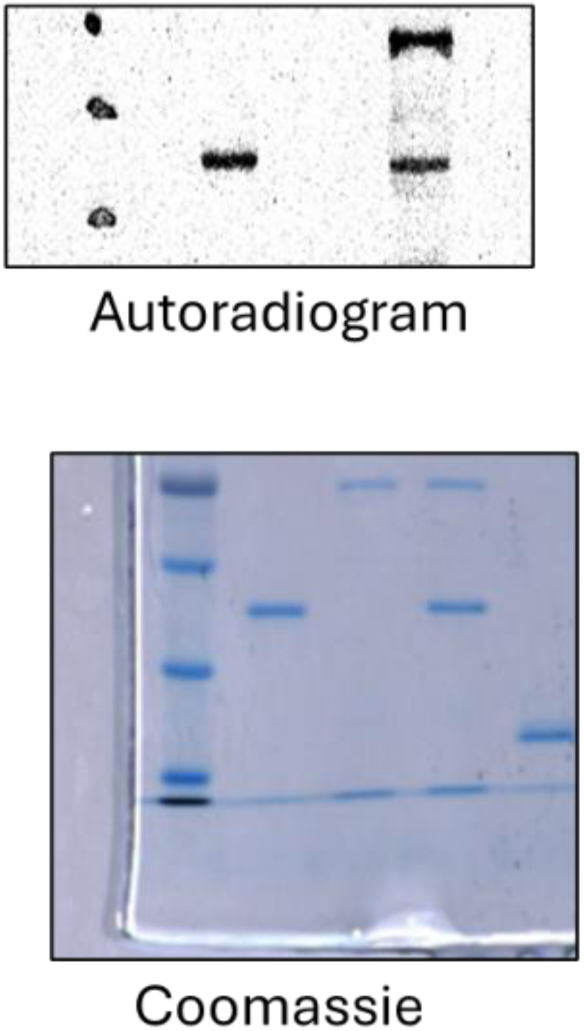
Original blots for Figure 2A

**Fig S6.**
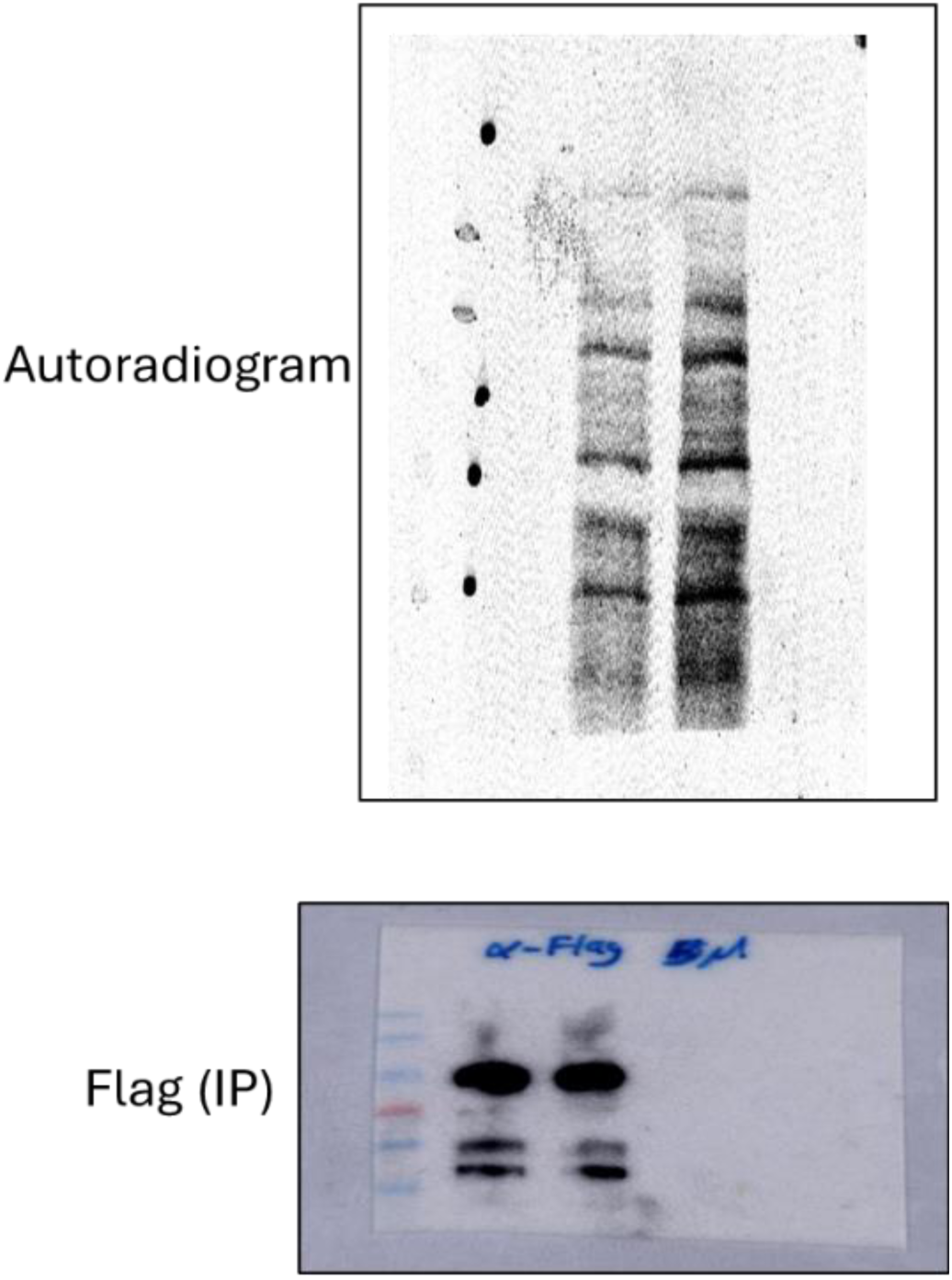
Original blots for Figure 2B

**Fig S6.**
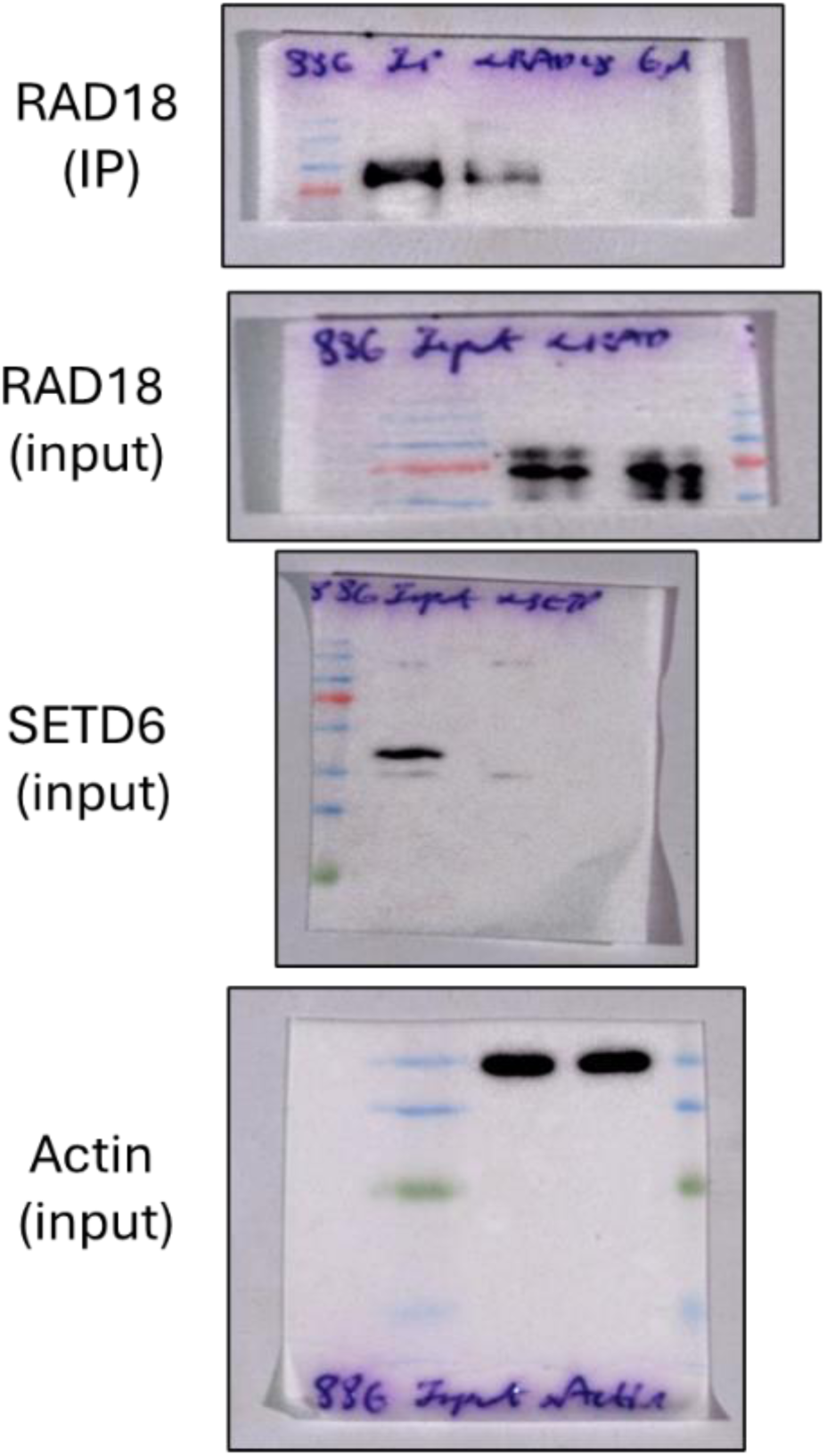
Original blots for Figure 2C

**Fig S7.**
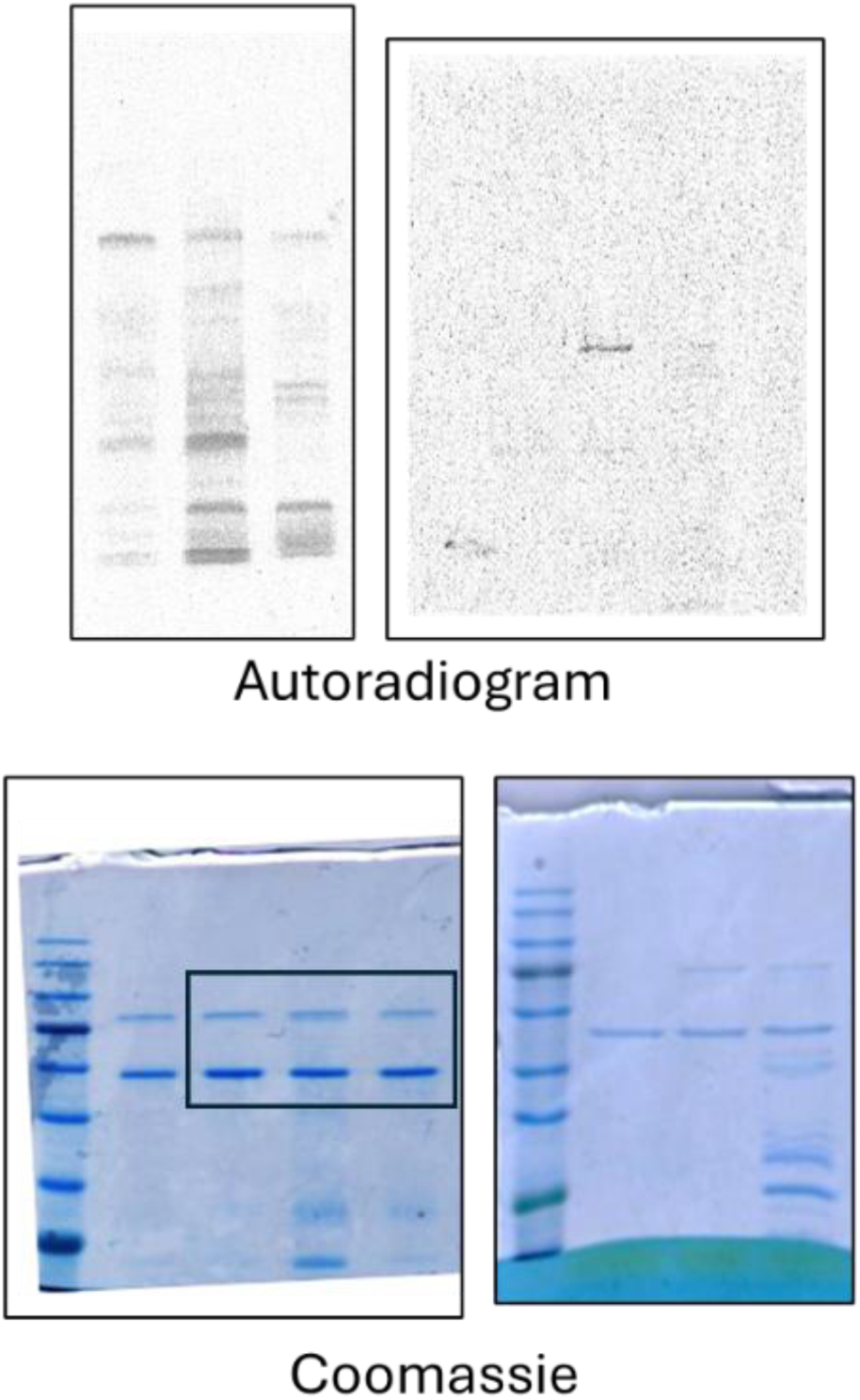
Originalblots for Figure 2E

**Fig S8.**
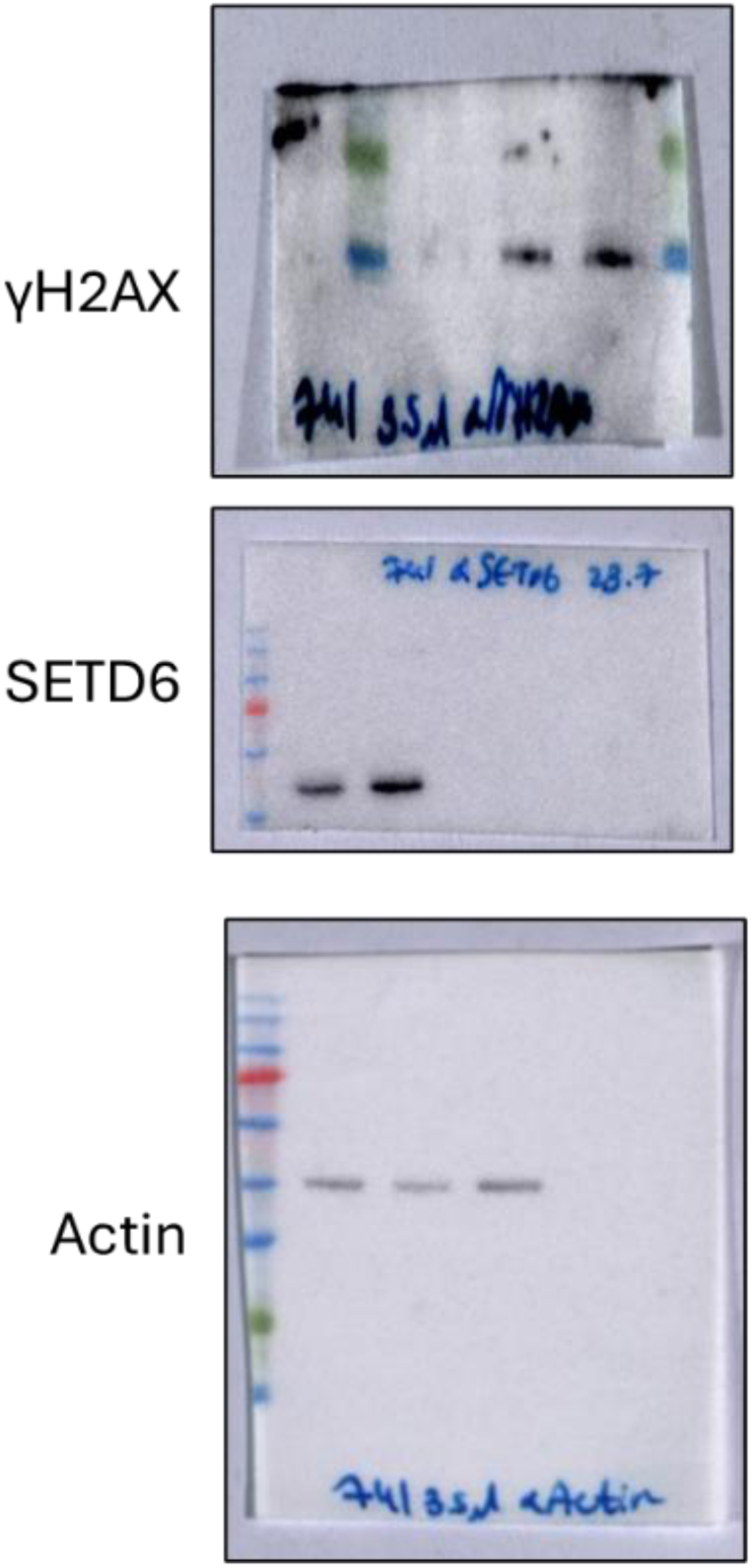
Original blots for Figure 4A

**Fig S9.**
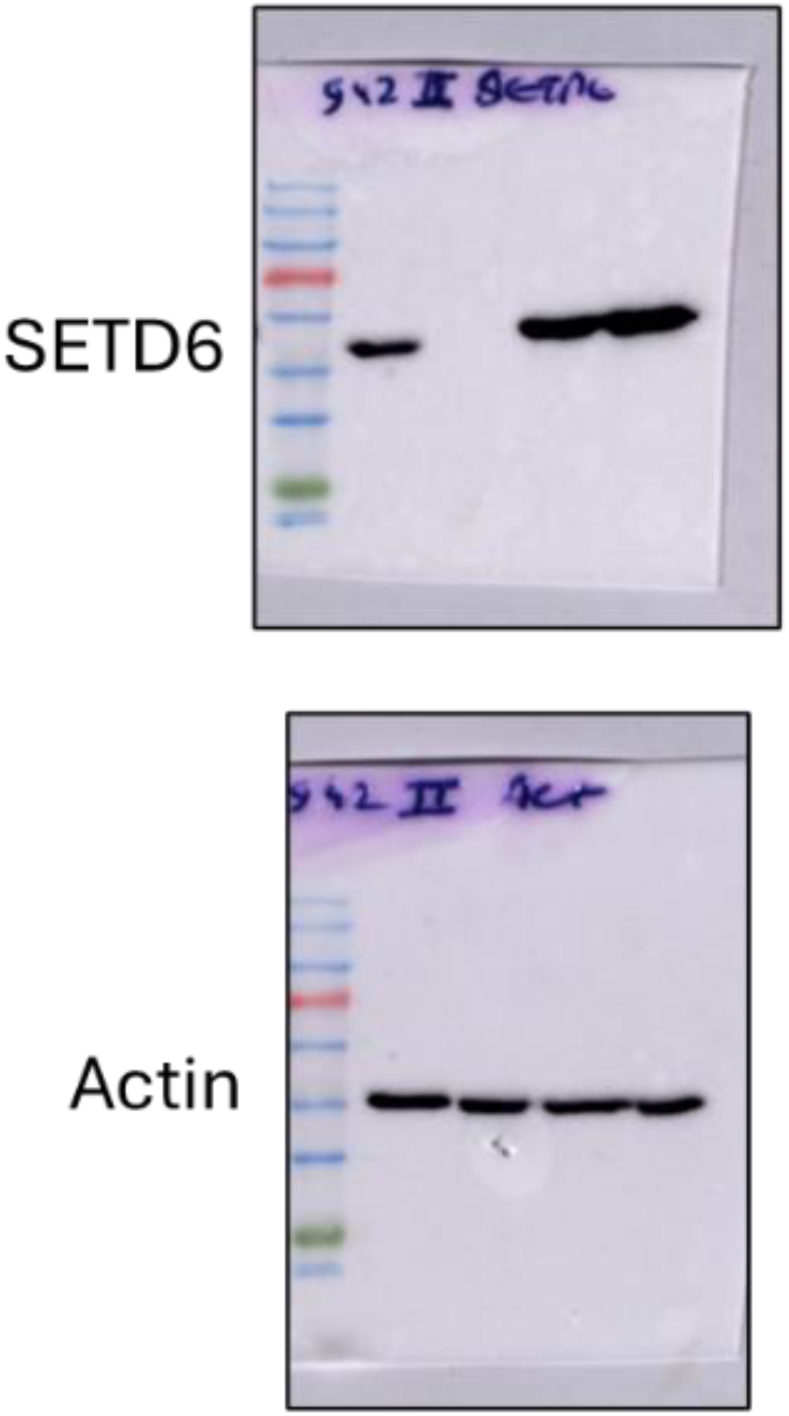
Original blots for Figure 4D

**Fig S10.**
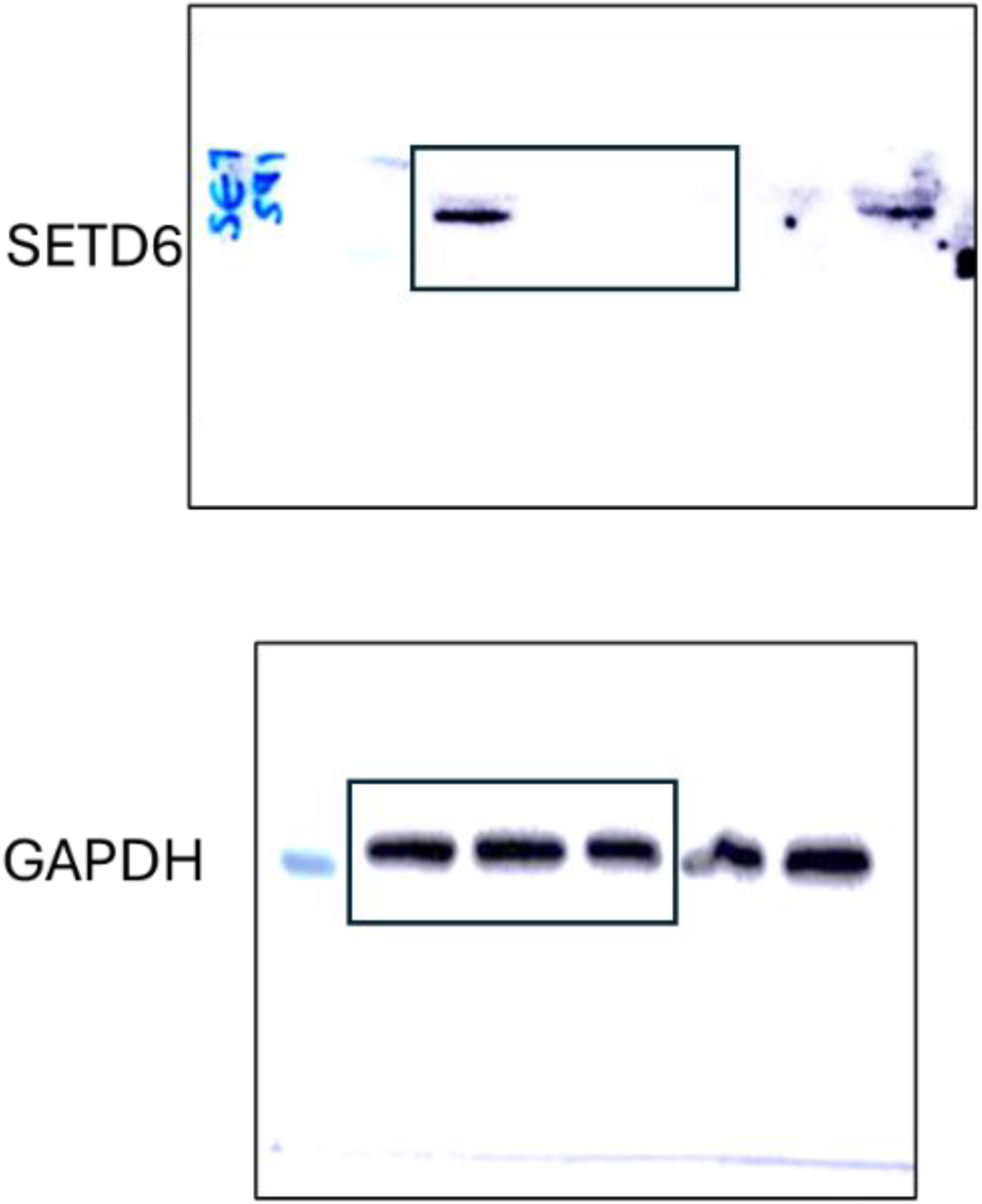
Original blots for Figure S3A

**Fig S11.**
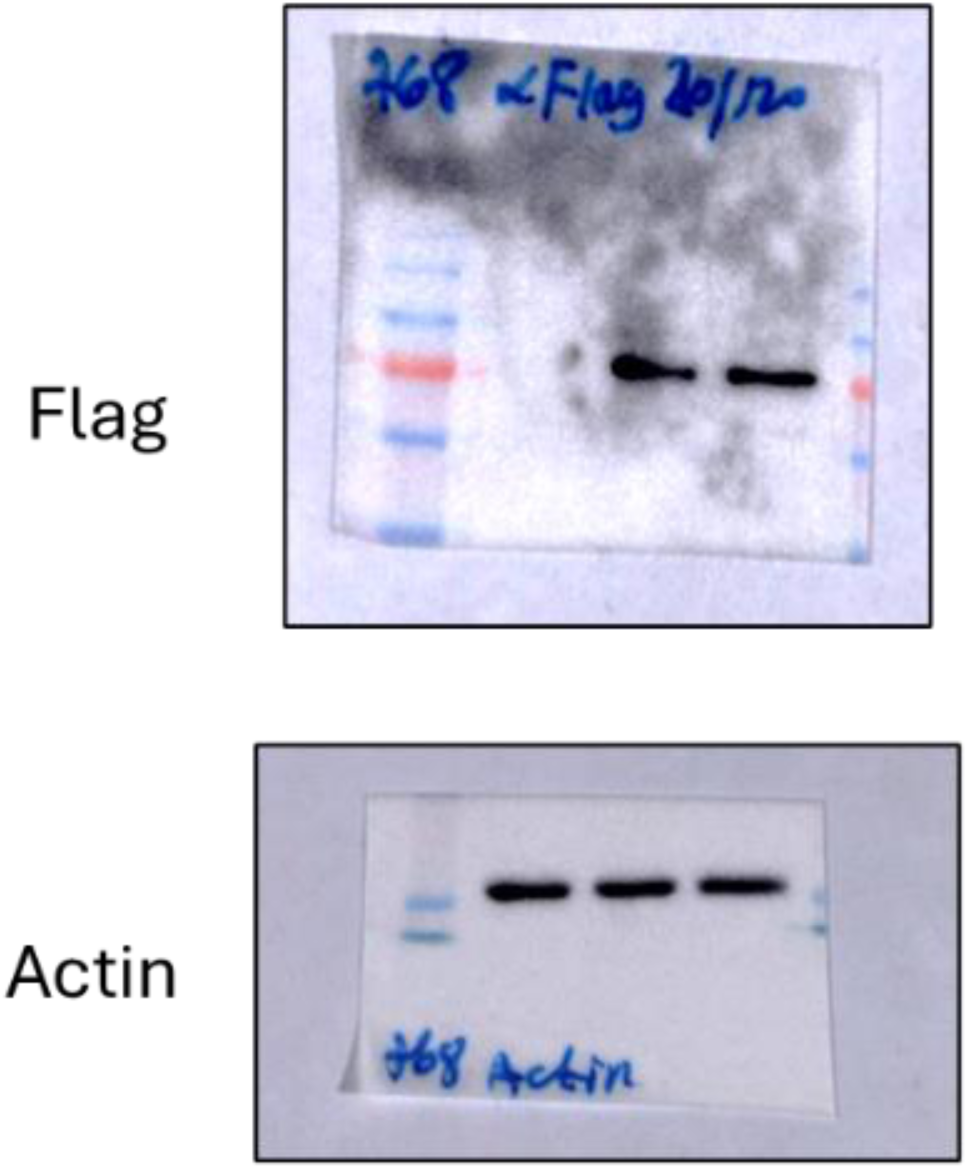
Original blots for Figure S3B

